# Sabatier principle for rationalizing enzymatic hydrolysis of a synthetic polyester

**DOI:** 10.1101/2022.03.30.486378

**Authors:** Jenny Arnling Bååth, Kenneth Jensen, Kim Borch, Peter Westh, Jeppe Kari

**Affiliations:** Department of Biotechnology and Biomedicine, Technical University of Denmark, Søltofts Plads, DK-2800 Kgs. Lyngby, Denmark; Novozymes A/S, Biologiens Vej 2, DK-2800 Kgs. Lyngby, Denmark; Department of Science and Environment, Roskilde University, Universitetsvej 1, DK-4000, Roskilde, Denmark

**Keywords:** PET hydrolase, polyester degradation, enzyme kinetics, heterogeneous catalysis, Sabatier principle, volcano curve, enzyme affinity

## Abstract

Interfacial enzyme reactions are common in nature and in industrial settings, including the enzymatic deconstruction of poly(ethylene terephthalate) (PET) waste. Kinetic descriptions of PET hydrolases are necessary for both comparative analyses, discussions of structure-function relations and rational optimization of technical processes. We investigated whether the Sabatier principle could be used for this purpose. Specifically, we compared the kinetics of two well-known PET hydrolases, leaf-branch compost cutinase (LCC) and a cutinase from the bacterium *T. fusca* (TfC) when adding different concentrations of the surfactant cetyltrimethylammonium bromide (CTAB). We found that CTAB consistently lowered the strength of enzyme-PET interactions, while its effect on enzymatic turnover was strongly biphasic. Thus, at gradually increasing CTAB concentrations, turnover was initially promoted and subsequently suppressed. This correlation with maximal turnover at an intermediate binding strength is in accordance with the Sabatier principle. One consequence of these results is that both enzymes had too strong intrinsic interaction with PET for optimal turnover, especially TfC, which showed a 20-fold improvement of *k_cat_* at the maximum. LCC on the other hand had an intrinsic substrate affinity closer to the Sabatier optimum and the turnover rate was 5-fold improved at weakened substrate binding. Our results show that the Sabatier principle may indeed rationalize enzymatic PET degradation and support process optimization. Finally, we suggest that future discovery efforts should consider enzymes with weakened substrate binding, since strong adsorption seems to limit their catalytic performance.

**ToC graphics:** 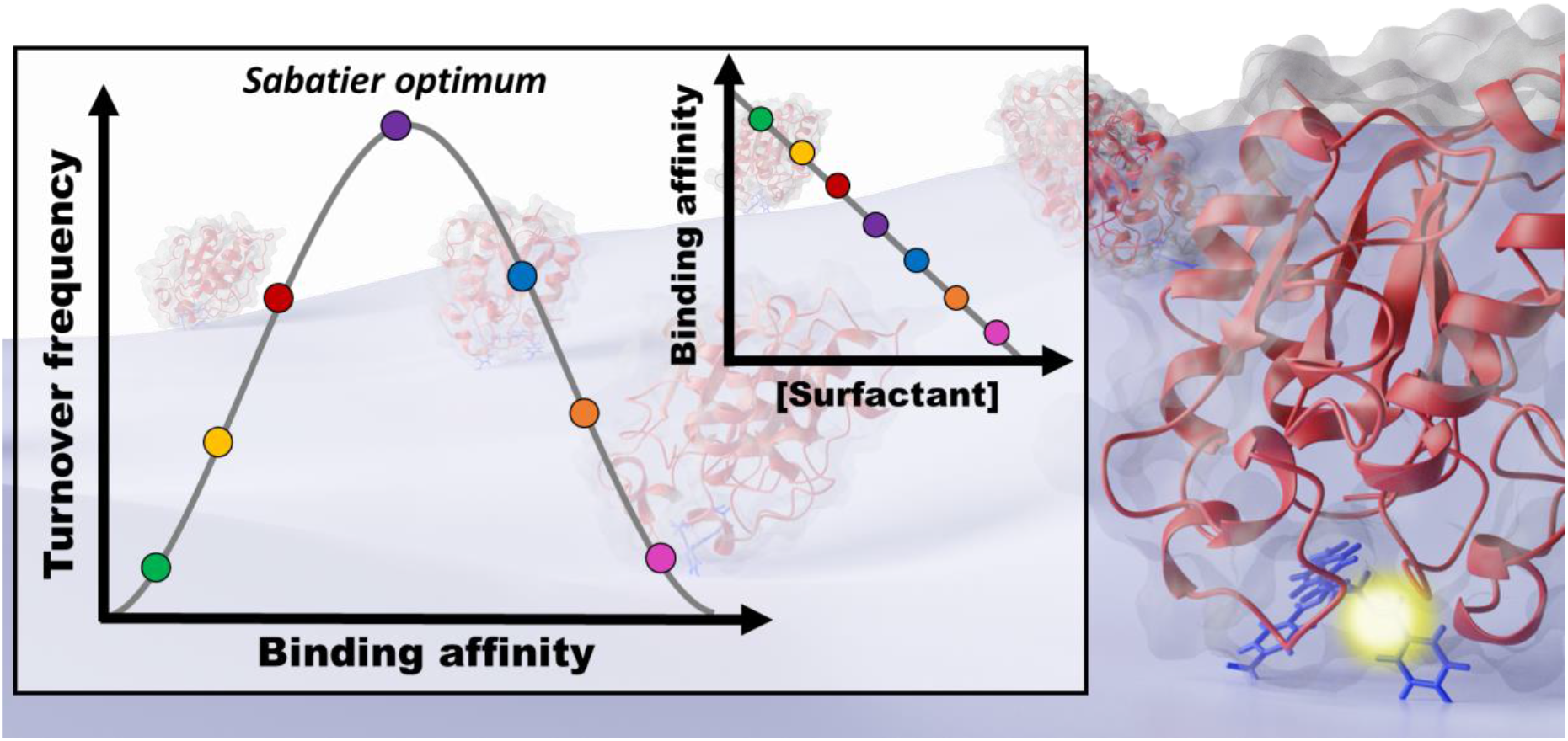

## Introduction

Enzymatic hydrolysis of poly(ethylene terephthalate) (PET) has been recognized as a potential technology for bioprocessing of some plastic waste streams.^1^ Several studies have reported promising enzyme candidates, including the PETase from *Ideonella sakaiensis,^2^* the leaf-branch compost cutinase LCC^3,4^ and cutinases from various *Thermobifida* species.^5–8^ However, there is still a need to engineer better catalytic efficiency and improve reaction conditions on PET; a substrate the enzymes were not evolved to act on. Previous activity in this area has used strategies like rational re-design of the active site,^9,10^ construction of chimeric enzymes with binding modules^11,12^ and application of surfactants in the reaction medium.^13,14^

To optimize enzymes rationally, it is crucial to have a molecular-level understanding of structure-function relationships. The experimental input that links structure and function is typically kinetic data, but there is no general framework to rationalize the kinetics of these interfacial enzymes, and this has hampered fundamental and comparative descriptions of PET hydrolases. In lack of a detailed kinetic model, the PET hydrolase reaction may be coarsely described by its most basic steps; complexation, catalysis, and dissociation, as shown in Figure 1A. In this three-step model, the first involves enzyme adsorption on the substrate surface and transfer of a piece of the PET molecule into the enzyme’s active site. The second step encompasses the covalent changes of the hydrolytic reaction (catalysis), while the third delineates product release and the enzyme’s dissociation to the aqueous bulk. This model bears some resemblance to the classical Henri-Michaelis-Menten scheme, but as the substrate is insoluble, the complexation step differs fundamentally. Unlike enzymes acting in the bulk phase, interfacial enzymes such as PET hydrolases need to compete with attractive forces in the substrate matrix to make a productive enzyme-substrate complex^15–17^ (see Figure 1B). One consequence of this is that the effective substrate concentration in interfacial enzyme reactions may be correlated with ligand binding strength. Thus, the tighter the binding, the more diverse polymer conformations on the surface could potentially be transferred to the active site.^18–20^ However, high ligand binding affinity may come at the cost of turnover speed.^21^ This type of tradeoff between binding strength and rate is well-known within inorganic, heterogeneous catalysis, and was originally coined as the Sabatier principle, which states that efficient catalysis occurs when the catalyst binds its reactant with intermediate strength.^22,23^ The Sabatier principle has recently proven useful for predicting and rationalizing catalytic properties of cellulases^24,25^ and as a guide for computer-aided enzyme design and discovery.^26^

**Figure 1.**
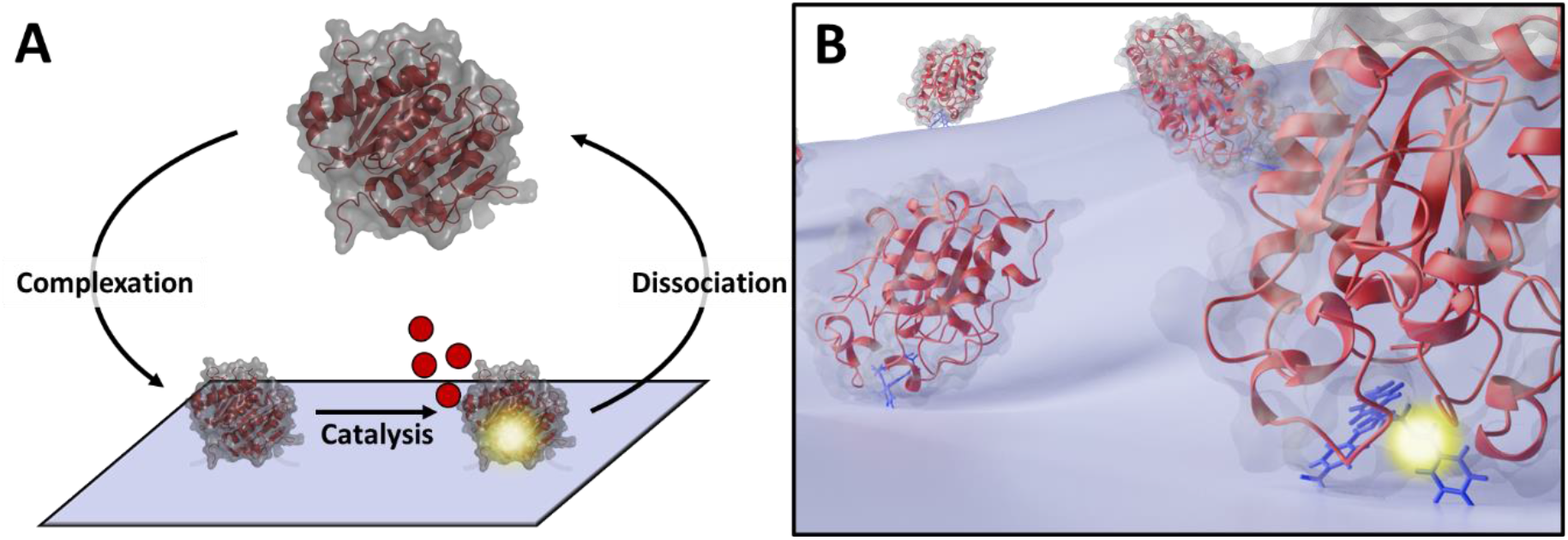
A) Simplified reaction scheme for enzymatic degradation of poly(ethylene terephthalate) (PET). PET hydrolase (red cartoon) binds to the PET surface and catalyzes the hydrolysis of ester bond(s), releasing soluble products (red disks), before enzyme and products dissociate into the solution. B) Illustration of the enzyme-substrate complex at the solid-liquid interface. The enzyme needs to dislodge a small piece of the polymeric substrate from the matrix to access the scissile bond and make a productive enzyme-substrate complex.

In this study, we investigate whether the Sabatier principle could rationalize the kinetics of two well-known PET hydrolases; leaf-branch compost cutinase (LCC) and a cutinase from the thermophilic bacterium *T. fusca* (TfC). We used two types of steady-state measurements to assess kinetic parameters for the hydrolysis of suspended PET particles, and tuned enzyme-substrate binding strength through the addition of the cationic surfactant cetyltrimethylammonium bromide (CTAB).

## Results

We have applied a modified Michaelis-Menten (MM) approach to assess the kinetics of two PET hydrolases, LCC and TfC. Specifically, we collected one set of data in the conventional way with substrate excess (^conv^MM analysis), while another was made in the inverse concentration regime with enzyme excess (^inv^MM analysis). We characterized both enzymes at different concentrations of the cationic surfactant CTAB, and derived kinetic parameters for both ^conv^MM and ^inv^MM by non-linear regression as described in the Experimental section and in the Supporting Information (SI). Experimental data (circles) and results of the regression analyses (lines) are illustrated in Figure 2.

**Figure 2.**
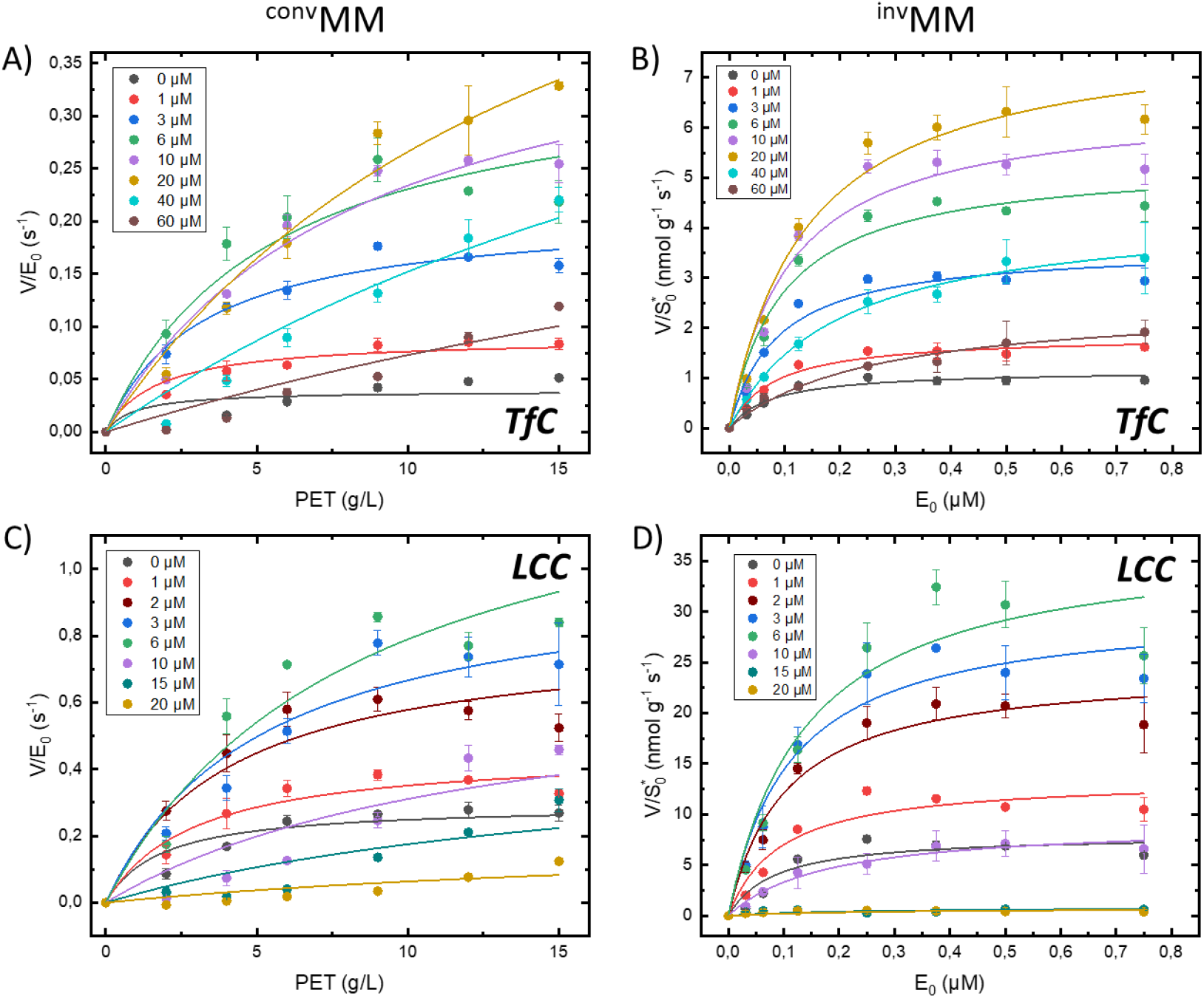
Experimental data at 50°C for conventional (panel A and C) and inverse (panel B and D) Michaelis-Menten analysis. Measurements (circles) were conducted for both LCC and TfC in buffers supplemented with different concentrations of the cationic surfactant CTAB. The CTAB concentrations are indicated by color codes in each panel. The results were analyzed with respect to the conventional- and inverse MM equation, eqs. (3) and (5), seen in the Experimental section. Error bars represent standard deviations of triplicate measurements.

Experimental values for the kinetic parameters *^inv^K_M_, ^inv^V_max_, ^conv^K_M_* and *^conv^V_max_* (defined in eqs. 2-6, Experimental section) derived in Figure 2, are listed in Table S1 in the SI. Inspection of these numbers revealed systematic effects of the surfactant. In particular, we found that the maximal specific rate from both the conventional (*^conv^V_max_*/E_0_, eq. 6) and inverse (*^inv^V_max_*/S*_0_, eq. 4) analyses showed distinctive maxima at intermediate CTAB concentrations (Figure 3). To put these observations into perspective, we reiterate the meaning of the two parameters. *^conv^V_max_*/E_0_ is the usual maximal turnover (eq. 6), which is experimentally observed when essentially all enzyme is complexed (when [ES]~E_0_). The inverse parameter, *^inv^V_max_*/S*_0_, specifies the rate when all attack sites on the PET surface is covered with enzyme (eq. 4). The effect of CTAB on these two parameters was distinctive and quite similar for both enzymes. Thus, for LCC both parameters increased approximately 5-fold between [CTAB] = 0 and the maximum at [CTAB] ~ 6μM (Figure 3). The analogous increment for TfC was up to 20-fold, and it follows that, CTAB strongly lessened the difference in performance of a highly efficient (LCC) and a mediocre (TfC) PET hydrolase.

**Figure 3.**
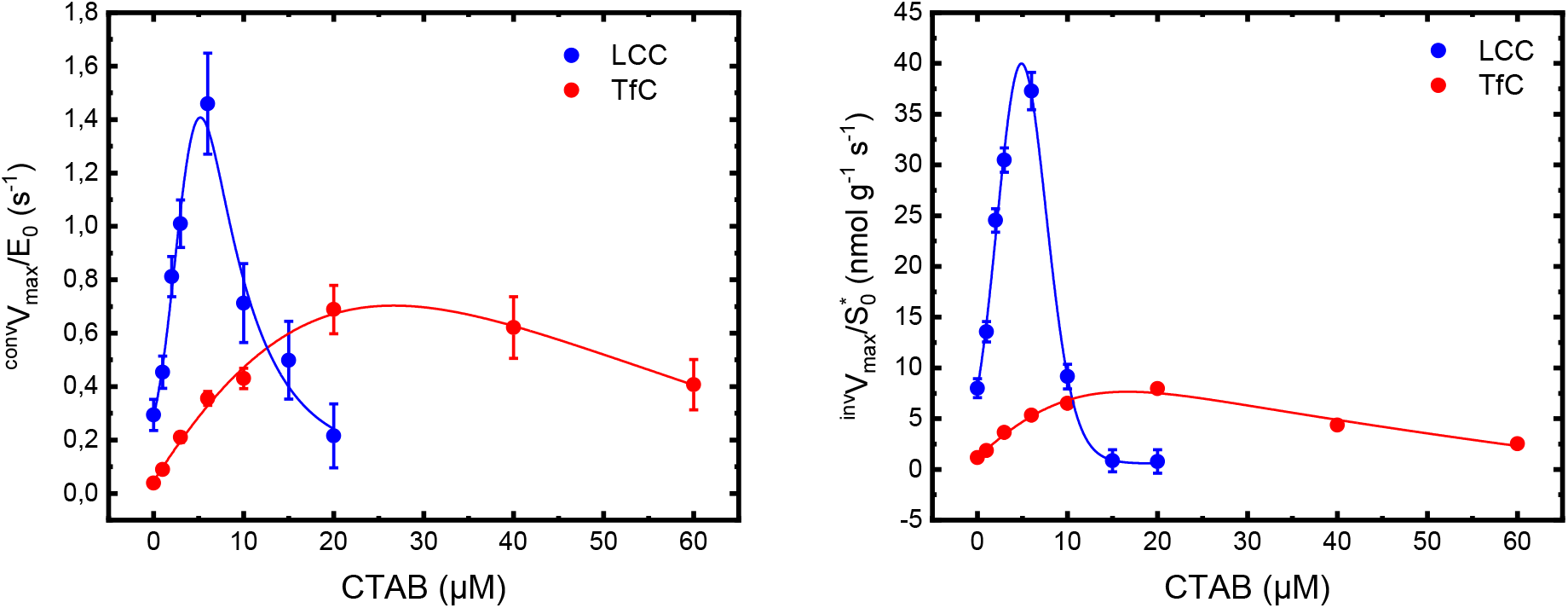
Maximal specific rates for conventional (^conv^V_max_/E_0_) and inverse (^inv^V_max_/S*_0_) MM analysis of LCC and TfC acting on PET particles at 50 °C. Symbols represent kinetic parameters obtained with different concentrations of the cationic surfactant CTAB (see Figure 2). Maximal specific rates from the two analyses showed distinctive optima at intermediate CTAB concentrations for both enzymes.

We conducted several experiments to explore possible origins of the biphasic curves in Figure 3. First, we found that at the concentrations used here, CTAB had no significant effect on the two enzymes’ thermostability, expressed as the apparent transition temperature, T_m_, (Figure 4A). In addition, we only found minor effects of CTAB on the catalytic performance against the small soluble substrate, 4-nitrophenyl butyrate (*p*NP-Bu). Specifically, when adding a (low) CTAB concentration corresponding to the ascending parts in Figure 3, we found no effect on the activity of LCC and a moderate inhibitory effect on TfC (Figure 4B). Hence, the large initial increments in Figure 3 could not be related to an intrinsic boost in enzyme activity generated by the surfactant. Finally, we assessed whether long-term exposure of the enzymes to high concentrations of CTAB, corresponding to the descending parts in Figure 3, resulted in a subsequent decrease in kinetic performance towards *p*NP-Bu. The results (Figure S3 in the SI) did not reveal any loss of enzyme activity, and we conclude that CTAB does not significantly influence the kinetic stability of the enzymes.

**Figure 4.**
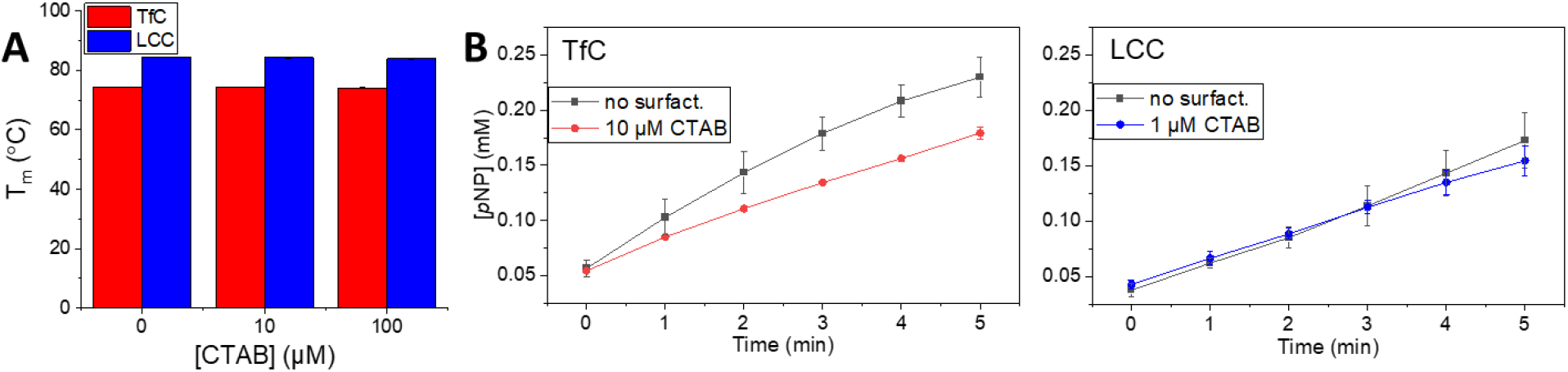
A) The apparent transition temperature for LCC and TfC in the absence and presence of CTAB at 10 or 100 μM. B) Progress curves for LCC and TfC acting on the soluble substrate pNP-Bu at no or a low concentration of CTAB. Experiments were conducted in duplicates and standard errors represent the spread. CTAB had no considerable effect on either the thermostability of the enzymes or their catalytic performance on the small soluble pNP-substrate.

Figure 5 illustrates adsorption behavior of LCC and TfC on the insoluble PET substrate (in the absence of CTAB). We analyzed these measurements with respect to a simple Langmuir isotherm

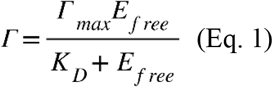

where Γ and Γ_max_ are respectively coverage and saturation coverage in mol enzyme per g substrate, E_free_ is the molar concentration of free enzyme (unbound enzyme) and *K_d_* is the dissociation constant. Lines in Figure 5 represent the best fits of eq. (1), and we found that TfC adsorbed more tightly (*K_d_*=22 ± 1.9 nM) compared to LCC (*K_d_*=260 ± 54 nM). However, saturation coverage was higher for LCC, 40 ± 4.1 nmol/g PET, compared to 15 ± 0.025 nmol/g for TfC.

**Figure 5.**
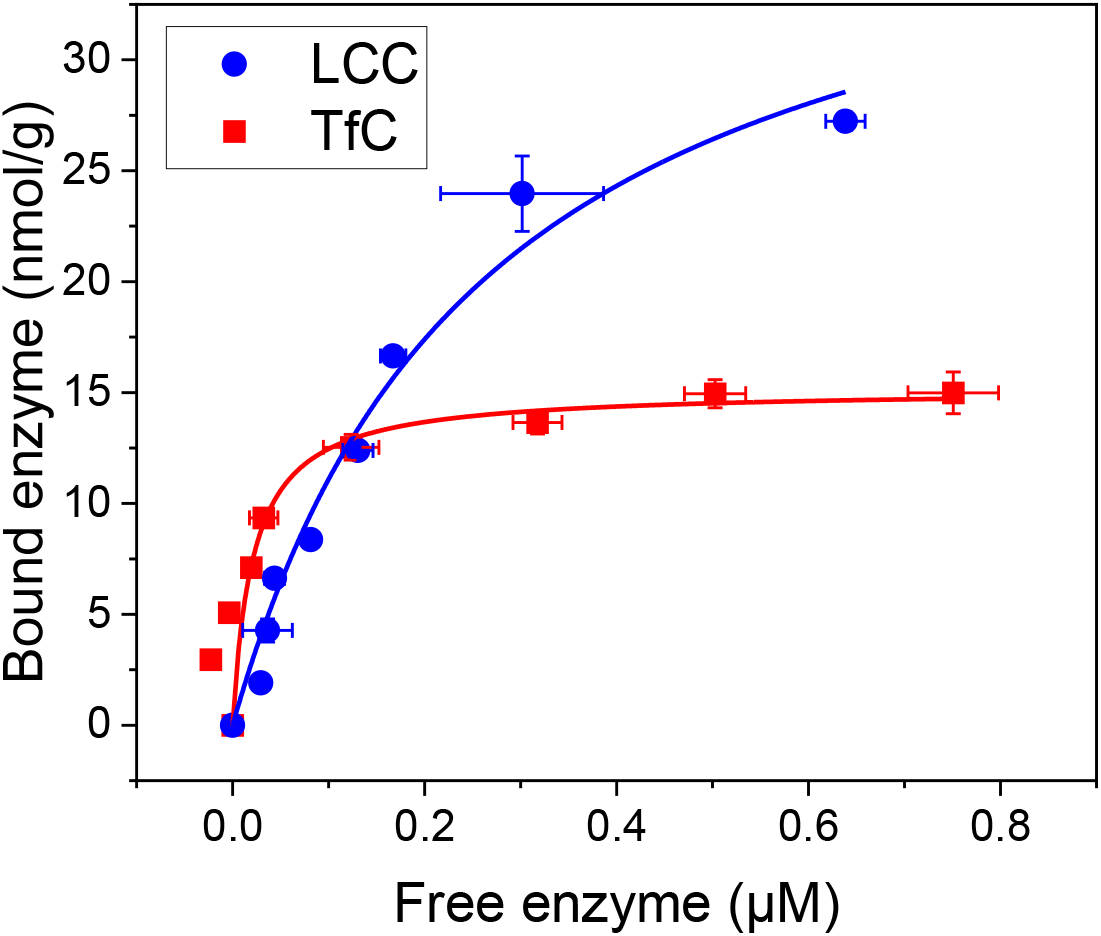
Enzyme adsorption on suspended PET particles (50 g/L) at 50 °C. Symbols represent the measured coverage, Γ, as a function of the free enzyme concentration, and lines are best fits of eq. (1). Error bars represent standard deviations of triplicate measurements. TfC showed the tightest adsorption, whereas saturation coverage was substantially higher for LCC, when comparing the two enzymes.

## Discussion

In recent years, gradually better PET degrading enzymes have been identified, and this has spurred optimism regarding bioprocessing of PET waste.^27–31^ However, continuous improvement of catalytic efficiency through engineering and discovery of novel enzymes will probably be necessary to make biological approaches viable contributions to a circular plastic economy. Progress in this field would benefit from a general framework to compare and rationalize the performance of different enzymes and conditions, and in the current work, we propose one potential approach to this. The two main elements in the suggested framework are a modified Michaelis-Menten analysis (Figure 2) and the Sabatier principle.

Our results (see Figure 3) revealed that addition of the surfactant CTAB had a distinctive and bi-phasic effect on the catalytic performance of two PET hydrolases. Especially for TfC, moderate amounts of CTAB (20-30 μM) substantially accelerated the maximal rate for both conventional and inverse MM kinetics. This CTAB concentration is one or two orders of magnitude below the critical micelle concentration,^32^ and it did not exert any measurable reduction in the thermostability of the investigated enzymes (Figure 4A). Neither did it significantly alter the catalytic efficacy on a soluble substrate (Figures 4B and S3), and in light of this, we will test whether the biphasic behavior could reflect CTAB-induced alterations in the strength of interactions with the insoluble substrate.

This idea has been discussed before. In particular, Furukawa and co-workers reported a beneficial effect of charged surfactants on the enzymatic hydrolysis of PET films. They found that an anionic surfactant promoted activity of the positively charged PETase from *Ideonella sakaiensis*,^14^ and that a cationic surfactant boosted activity of another PET hydrolase, which carried a net-negative charge at the experimental pH.^13^ These observations were collectively ascribed to enhanced enzyme adsorption driven by electrostatic interactions between enzyme and an oppositely charged surfactant that accumulated on the PET surface. However, this interpretation is not readily transferred to the current data. Firstly, we observed both activating and inhibitory effects of the same surfactant (Figure 3). Secondly, the two investigated enzymes have pI values respectively above (LCC: pI 9.3) and below (TfC: pI 6.3) the experimental pH (pH 8.0). Nevertheless, the two enzymes responded analogously to the addition of CTAB, and the concomitant buildup of positive charge on the PET surface.

In search of a more robust interpretation of the biphasic behavior in Figure 3, we note that the Michaelis constant may be used as a descriptor of substrate affinity.^33,34^ Thus, while not a true binding constant, the concentration required to reach half-saturation under different conditions provides some ranking of substrate interaction strength. As illustrated in Table S1A in the SI, we found that K_M_ values increased regularly with the surfactant load throughout the investigated range. We conclude that CTAB consistently lowered the strength of enzyme-substrate interactions and we will discuss the results in this perspective. We acknowledge that other more specific effects of CTAB may influence the maximal specific rates, and we will return to this in the concluding paragraph.

General relationships between binding strength and catalytic turnover can be expressed by the Sabatier principle,^35^ which states that catalysis is most effective when the interaction between catalyst and reactants is intermediate in strength. The intuitive underpinning is that tight binding implies slow dissociation of stable enzyme-substrate intermediates, while weak binding is associated with low complex concentration. Both of these limiting cases lead to a poor overall rate, while an intermediate binding strength balances off the two effects and hence supports a faster turnover. The Sabatier principle has been widely employed within inorganic, heterogeneous catalysis,^36^ but it has also been applied to interfacial enzyme reactions.^24,37^ Experimentally, the principle can be illustrated in so-called volcano plots, which have a measure of catalytic efficacy such as turnover frequency on the ordinate and interaction strength on the abscissa. For the current systems, this implies plotting the maximal specific rate from either ^inv^MM or ^conv^MM as a function of binding strength expressed as a Michaelis constant. Figure 6 illustrates such plots, where we used *^inv^K_M_* (i.e. the enzyme concentration required to reach half-saturation under conditions of enzyme excess) as a descriptor of binding strength. These plots clearly had volcano-like shapes, and hence corroborated that intermediate binding strength provides the most efficient catalysis in accord with the Sabatier principle.

**Figure 6.**
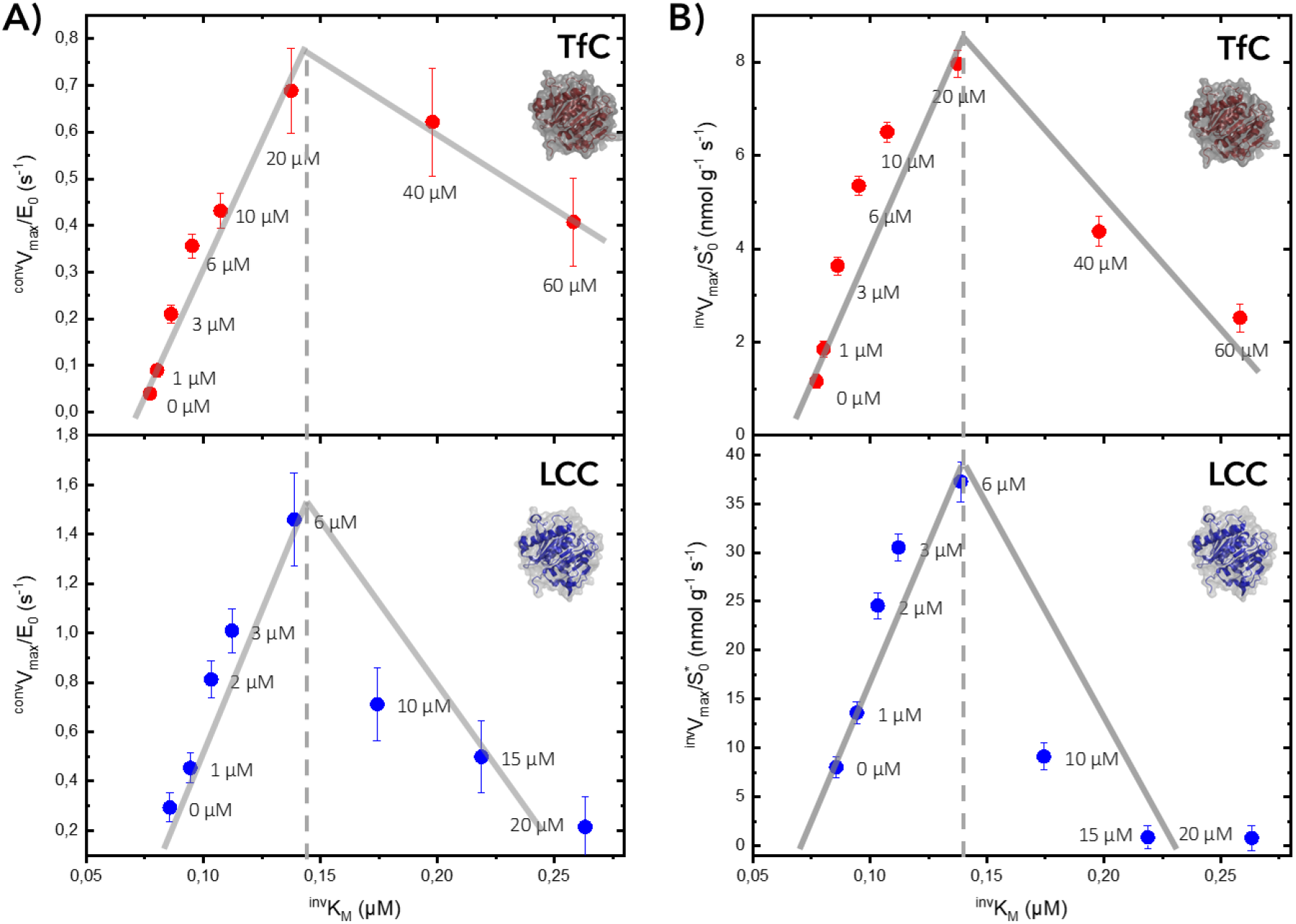
Volcano plots for TfC and LCC acting on insoluble PET. The turnover frequency under A) enzyme saturation condition (^conv^V_max_/E_o_) or B) substrate saturation condition (^inv^V_max_/S*_o_) is plotted against the enzyme-substrate binding affinity (^inv^K_M_) for different concentrations of CTAB in the background, as specified by the label in the plot. Symbols represent kinetic parameters, and solid lines are to guide the eye.

One consequence of data in Figure 6 is that LCC and particularly TfC bind their substrate too tightly for efficient catalysis. The tight binding of TfC was reflected both in a low *^inv^K_M_* (77 ± 15nM, Table S1) and the independently measured *K_d_* value (22 ±1.9 nM, Figure 5). This strong substrate affinity occurred together with slow maximal turnover of TfC (*k_cat_* ~ 0.04 s^-1^), but when the interaction was weakened by CTAB to a level of *^inv^K_M_* ~ 140 nM, the turnover rose dramatically to about 0.8 s^-1^. Interestingly, an *^inv^K_M_* of this magnitude also gave rise to the highest values of both *k_cat_* for LCC (1.6 s^-1^, Figure 6A) and maxima in the inverse specific rates for both enzymes (Figure 6B). Hence, it appears that a substrate affinity specified by *^inv^K_M_* ~ 140 nM represents the Sabatier optimum, where the lifetime of the enzyme-substrate complex attains a favorable, intermediate value. As the inherent substrate affinity measured without CTAB (see Figure 5) was lower for LCC compared to TfC, it required less CTAB to bring LCC to the Sabatier optimum (Figure 6). This implied that the inherent substrate affinity of LCC was closer to the Sabatier optimum, which explains the larger activity increment (Figure 6A) at the maximum of TfC (20-fold) compared to LCC (5-fold). The binding affinity of LCC appears to be better tuned for the substrate, and hence there is less to gain for this enzyme.

Comparisons of LCC and TfC may be expanded by considering specific changes in respectively ^inv^MM and ^conv^MM parameters. To this end, we already noted that the maximal turnover, *k_cat_* = *^conv^V_max_*/E_0_ (eq. 6) at the Sabatier optimum was about twice as high for LCC than for TfC (Figure 6A). Comparisons of the inverse maximal rate *^inv^V_max_*/S*_0_, illustrated in Figure 6B, are more complicated as it reflects the product of *k_cat_* and Γmax (eq. 4). From the binding isotherm in Figure 5 it can be seen that LCC had around two times more binding sites than TfC. As both *k_cat_* and Γ_max_ were 2-fold higher for LCC, we would expect that this enzyme performed 4-fold better than TfC under the most favorable conditions (*i.e*. when the substrate is saturated with enzyme and the binding strength is adjusted to the Sabatier optimum). This prediction is confirmed in Figure 6B, where direct and independent measurements of *^inv^V_max_*/S*_0_ gave maximal values of about 9 nmol g^-1^ s^-1^ and 40 nmol g^-1^ s^-1^ for TfC and LCC, respectively. This supports the conclusion that the superior performance of LCC on PET, which is widely recognized,^3,4,28,38^ reflects partly a low, nearly optimal substrate affinity (which is associated with a high *k_cat_*) and partly a high capacity of combining productively with different polymer conformations on the PET surface (high attack site density). If indeed so, it would be relevant to investigate other PET hydrolases with variable substrate affinity. So far, focus has been on high affinity,^11,12,39–43^ but the current work hints that this strategy may not always be fruitful. Hence, we suggest that future discovery campaigns consider enzymes with a broad spectrum of substrate binding strengths.

In conclusion, we have found that effects of the cationic surfactant CTAB on the kinetics of two PET-hydrolases may be rationalized along the lines of the Sabatier principle. We hasten to say, that other, more specific effects of the surfactant cannot be ruled out by the current experiments. However, controls focusing on thermodynamic- and kinetic enzyme stability as well as the general catalytic performance against soluble substrate failed to explain the pronounced kinetic alterations observed on insoluble PET. Instead, we propose that the biphasic effect of CTAB on the enzymatic turnover reflected a continuous weakening in enzyme-substrate interactions as surfactant concentrations rose. This weakening initially promoted and subsequently suppressed enzyme activity as stipulated by the Sabatier principle. If indeed applicable, the combination of the modified MM approach illustrated in Figure 2 and the empirical Sabatier principle may be a useful tool for rationalizing and optimizing enzymatic PET degradation. It may, for example, serve to elucidate structure-function relationships in engineered enzyme variants, as well as identify better reaction conditions for PET hydrolases.

## Experimental section

### Enzymes

Two cutinases, Leaf-branch compost cutinase (LCC) [PDB: 4EB0] and TfC from *Thermobifida fusca* [PDB: 5ZOA] were heterologously expressed in *Bacillus subtilis* and purified in a similar way as described previously.^44,45^ The two enzymes are of the same size (28 kDa) and have a sequence identity of approximately 60 %. The production of LCC incorporated the following two modifications compared to the published procedures. The native signal peptide was replaced by the signal peptide from the *B. licheniformis* α-amylase AmyL (FJ556804.1), and a histidine tag (6xH) was added to the C-terminal. A small linker consisting of LE was inserted between the C-terminal and His-tag. The fermentation broth was sterile filtrated and 500 mM NaCl was added and adjusted to pH 7.5/NaOH. The sample was loaded onto a Ni-Sepharose™ 6 Fast Flow column (GE Healthcare, Piscataway, NJ, USA) equilibrated in 50 mM HEPES, pH 7.5 with 500 mM NaCl (buffer A). After loading, the column was washed with 10 column volumes of buffer A, and bound proteins were eluted with 500 mM imidazole in buffer A. The fractions containing the enzyme were pooled and applied to a Sephadex™ G-25 (medium) (GE Healthcare, Piscataway, NJ, USA) column equilibrated and eluted in 50 mM HEPES pH 7.5. Enzyme concentrations were determined by Abs280. Molar extinction coefficients and theoretical pI values were calculated using the protein identification and analysis tools in the ExPASy Server.^46^ T_m_ of the enzymes was determined by differential scanning fluorimetry, using a Nanotemper Prometheus Nt.48 (Nanotemper). Enzyme samples (in pure phosphate buffer or supplemented with 10 or 100 μM CTAB) were heated from 20 °C to 95 °C at 10 % laser intensity and a rate of 1.5 °C/min.

### Substrates and chemicals

The PET substrate used was a semi-crystalline PET powder purchased from Goodfellow Co. UK (Product number ES306031). The typical particle size was about 100 μm.^47^ The powder was suspended in 50 mM sodium phosphate (NaPi) buffer pH 8.0. The surfactant Cetyltrimethylammonium bromide (CTAB; 57-09-0), and the substrate 4-Nitrophenyl butyrate (*p*NP-Bu; 2635-84-9) were both purchased from Sigma.

### Binding isotherms

The adsorption of TfC and LCC to PET was determined using a PET concentration of 50 g/L and (total) enzyme concentrations ranging from 0-2 μM. Adsorption measurements used 1-hour equilibration in a thermomixer operated at 1000 rpm at 50 °C, and non-binding microtiter plates (Greiner Bio-One™) were used to reduce unproductive binding. Solids and liquids were separated by centrifugation in a temperature-controlled centrifuge, set to 50 °C. The protein content of the supernatant was determined using a micro BCA protein kit from Thermo Fischer scientific (Product number 23225), as described previously.^47^ Standard curves of the two enzymes (ranging from 0-2 μM in concentration) were used to quantify the amount of free enzyme in the reactions (performed in triplicates). The bound enzyme population was calculated from the difference between the total and the free concentrations, and the bound fraction was used to assess binding parameters as described previously.^47^

### Activity assay with pNP-Bu

To investigate potential side effects of the surfactant CTAB on the two enzymes, not related to surface phenomena and the Sabatier principle, we investigated activity on a soluble *p*NP-substrate at different concentrations of CTAB. One set of experiments was performed in order to investigate if long contact time with high amount of CTAB present (concentrations that resulted in a decrease in PET activity) possibly resulted in (irreversible) enzyme denaturation. This control was performed in two steps, with the first involving incubation of the enzymes with either CTAB or in pure 50 mM NaPi buffer at 50 °C, shaking at 300 rpm. The samples were incubated in microtiter plates with 10 μM enzyme and 0, 10 or 40 μM CTAB (LCC) or 0, 40 or 80 μM CTAB (TfC). After 2 hours, the samples were 1000-fold diluted to final enzyme concentrations of 10 nM. These enzyme dilutions were used in a second step, where enzyme activity was monitored on *p*NP-Bu. In a microtiter-plate, 5 nM enzyme and 5 mM *p*NP-Bu were mixed and the enzymatic release of *p*NP was monitored over 5 minutes in a plate reader at 405 nm at 25 °C. Reactions were performed in duplicates and blank samples without enzymes were included. The concentration of released *p*NP was calculated from standards with known concentrations.

We also assessed possible effects of CTAB on progress curves, which used the soluble *p*NP substrate. In a volume of 100 μL, reactions with 50 mM buffer, 5 nM enzyme, 5 mM *p*NP-Bu, either with or without CTAB (1 μM for LCC and 10 μM for TfC) were prepared and the enzymatic release of *p*NP was monitored and analyzed as explained above. The CTAB concentrations were selected to match concentrations resulting in an increased catalytic rate in the activity measurement on the insoluble PET substrate.

### Activity assay for solid PET substrate and Michaelis-Menten analysis

For determination of PET hydrolase activity on insoluble PET, a plate reader-based assay (Abs240) adapted for initial rate kinetics was used, described in detail elsewhere.^48^ Enzyme reactions were performed at 50 °C, in 50 mM NaPi buffer pH 8, using non-binding microtiter plates (Greiner Bio-One™), in an incubator operated at 450 rpm (KS 4000 ic control, IKA, Staufen, Germany). Initial activity measurements were performed in duplicates with a final volume of 250 μL, The load of PET was 10 g/L, enzyme concentrations were 0.10 μM and CTAB concentrations ranging from 0-100 μM. The contact time for these reactions were 2 hrs. for TfC and 30 min for LCC. Enzymatic product formation was quantified as “BHET-equivalents” (BHETeq), which were defined by the supernatant absorbance at 240 nm normalized against standard curves of BHET. Hence, the derived rates were based on soluble products only and this has previously been shown to be a good descriptor of the overall activity.^48^ Results from initial activity measurements are presented in Figure S4 in SI.

For kinetic analysis, two sets of experiments (each performed in triplicates) were executed, either under conditions of enzyme saturation (^conv^MM) or substrate saturation (^inv^MM). Experiments under ^conv^MM used 0.1 μM enzyme and a PET load from 0-20 g/L while ^inv^MM experiments used a fixed PET load of 10 g/L, and enzyme concentrations from 0-1 μM. Both types of experiments were conducted in pure buffer and in buffers supplemented by CTAB at concentrations ranging from 1-60 μM (TfC) and 1-20 μM (LCC). Other assay conditions were similar as explained above.

### Data analysis

The steady-state kinetics of LCC and TfC was analyzed under ^conv^MM and ^inv^MM conditions. The former used a constant and low enzyme concentration (E_0_) and recorded initial rates for a number of substrate loads (S*_0_, specified below). These results were analyzed by the conventional Michaelis-Menten equation (eq. 5). For ^inv^MM experiments, we used a fixed substrate load and measured initial rates at gradually increasing enzyme concentrations, E_0_. These latter data were analyzed using the inverse MM equation (eq. 3). Background, validity and limitations of this approach have been discussed in detail elsewhere,^49,50^ but we briefly reiterate the key relationships that are needed for the current discussion. The pivot of the analysis is the assumption that the mass-load of substrate (S*_0_) in g/L, which is known in kinetic experiments, can be converted to an apparent molar substrate concentration (S_0_) if one knows the density of accessible surface sites (in mol/g). This number can be approximated experimentally from a binding isotherm as the one shown in Figure 5, where the saturation coverage (Γ_max_) give the number of accessible surface sites on 1 g PET substrate. We note that Γ_max_ depends on both the enzyme and the physical properties (particularly surface area) of the insoluble substrate. The apparent molar concentration of substrate may be written

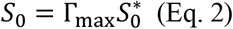

Under the condition of enzyme excess, the steady-state rate may be described using a rate equation, which is symmetric to the MM equation (eq. 5) and sometimes called the inverse MM equation.^50^

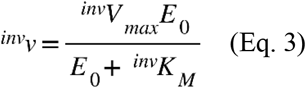

with *^inv^V_max_* being the maximal rate at substrate saturation (i.e. the rate when all accessible surface sites are in complex with an enzyme). This may be expressed as

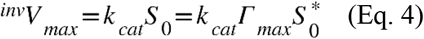

The Michaelis constant in eq. (3), *^inv^K_M_*, is the molar enzyme concentration at the half-saturation point. The conventional MM equation (eq. 5), which holds under conditions of substrate excess, may be expressed with *K_M_* in mass units (g/L) as discussed elsewhere.^50^

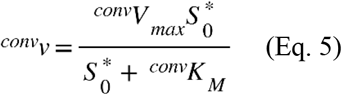

In eq. (5), *^conv^V_max_* is the maximal rate at enzyme saturation, defined in the usual way:

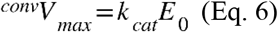

The Michaelis constant *^conv^K_M_* in eq. (5) has the unit of mass concentration (g/L), but can be converted to *^inv^K_M_* using the relationship *^inv^K_M_* = *^conv^K_M_Γ_max_*. We used the two MM equations (eqs. 3 and 5) to analyze the experimental data and the derived kinetic parameters to rationalize and compare the kinetics of LCC and TfC. Additional information on non-linear regression is described in detail in the SI.

## Supporting Information available

Detailed information on non-linear regression and global fit using competitive inhibition model with local *V_max_*, tables with extracted kinetic parameters, results from control experiments with high concentration of CTAB and initial activity measurements with increasing amount of CTAB.

## Acknowledgments

The research was supported by the Independent Research Fund Denmark (Grant number: 307 8022-00165B) and the Novo Nordisk foundation (Grant number: NNFSA170028392). The authors acknowledge Kristina Mielec for technical assistance with laboratory experiments.

## Notes

K.B. and K.J. work for Novozymes A/S, a major enzyme producing company.

