## Supporting Information for "Sabatier principle for rationalizing enzymatic hydrolysis of a synthetic polyester"

**Table of contents**

1. Non-linear regression and global fit using competitive inhibition model with local $V_{max}$

2. Control experiments with high concentration of CTAB

3. Initial activity measurements with increasing amount of CTAB

### 1. Non-linear regression and global fit using competitive inhibition model with local $V_{max}$

Initial fit to eqs. (3) and (5) (main text) showed somewhat linear increase in ${}^{conv}K_M$ and ${}^{inv}K_M$ for increasing concentration of CTAB (Figure S1), similar to what is seen for a competitive inhibitor. Thus, it was not possible to reach saturation (see Figure 2, main text) in the steady-state measurements with a high background of CTAB. As a consequence, the MM parameters showed a high parameter correlation at high CTAB concentration. This is illustrated in Figure S2, where the correlation between the MM parameters is plotted against the CTAB concentration. At high CTAB concentration the correlation approaches 1, since the experimental steady-state rates are far from reaching saturation (e.g. ${}^{conv}K_M > S_0$ or ${}^{inv}K_M > E_0$).

The relationship between the CTAB concentration (A) and the apparent $K_{M,app}$ could be accounted for using eq. (S1),

$$K_{M,app} = K_M\left(1 + \frac{A}{K_D}\right) \text{ (Eq. S1)}$$

where $K_d$ is the dissociation constant governing the interaction between CTAB and the enzyme (${}^{conv}$MM) or substrate (${}^{inv}$MM). To stear the fit, we substituted eq. (S1) in both eq. (3) and (5) and fitted the family of saturating curves using global non-linear regression with local $V_{max}$. We confirmed that this global regression approach did not infer a biased trend on the derived $K_{M,app}$ by comparing the global and local fit. In general, the best-fit parameters were similar between global and local fits, supporting that no significant bias was introduced in the global regression analysis (see Table S1). The global fit significantly improved the goodness of fit (as judged by the $R^2$, Table S1 ) and lowered the parameter dependency of the derived parameter (data not shown). For this reason, the global parameters were used in the data analysis. A full list of parameters from the two regression methods can be found in Table S1.

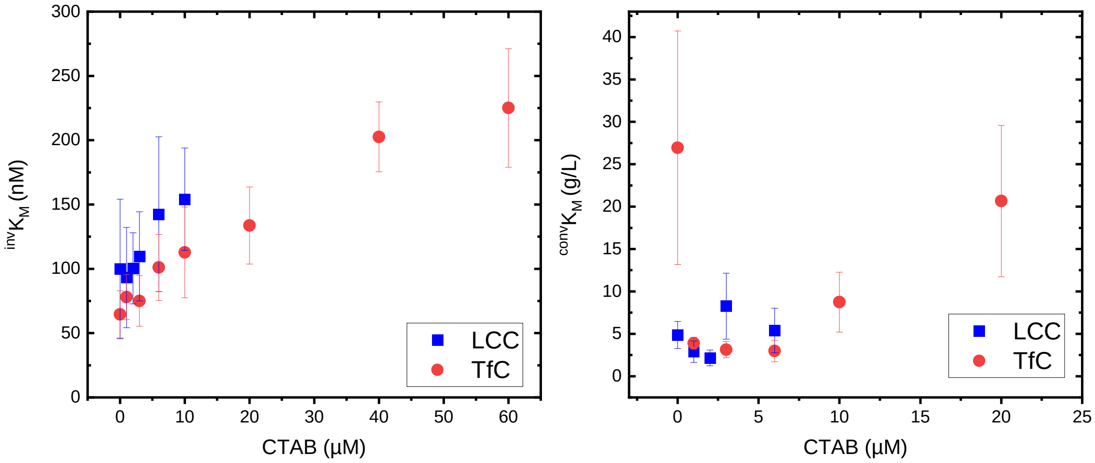


*Figure S1. Michaelis constants from local non-linear regression analysis to eqs. (3) and (5) (main text) as a function of CTAB concentration. Both* $^{inv}K_M$ *(left) and* $^{conv}K_M$ *(right) show a clear tendency to increase with CTAB concentration. At high CTAB concentration it was not possible to determine* $K_M$ *for LCC due to near linear MM curves.*

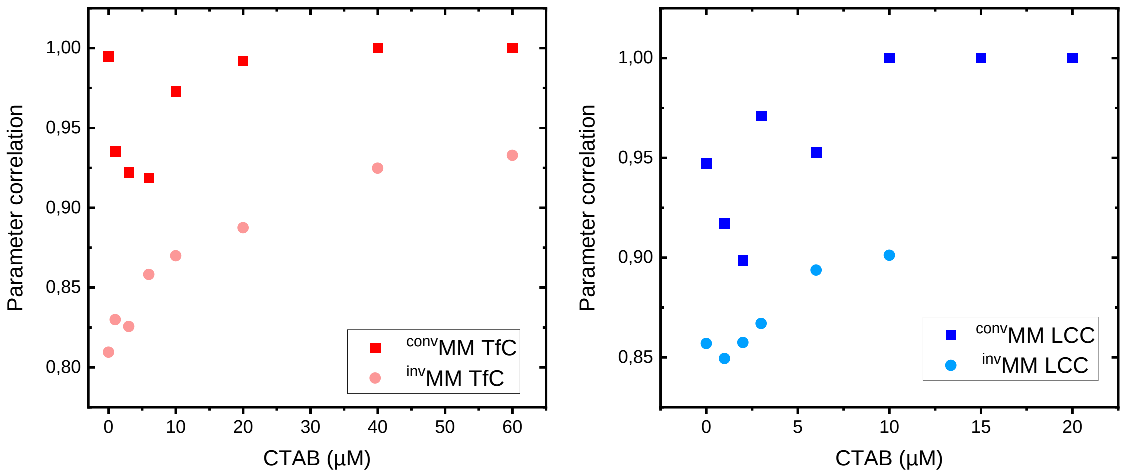


*Figure S2. Correlation between MM parameters from local regression analysis to eqs. (3) and (5) (main text). The derived* $^{conv}MM$ *and* $^{inv}MM$ *parameters showed a clear tendency for both TfC (left) and LCC (right). With increasing concentration of CTAB, the two parameters were highly correlated making it difficult to deconvolute them.*

**Table S1A. Parameters from global non-linear regression with local $V_{max}$**

| Enzyme | CTAB (μM) | $^{conv}V_{max}/E_0$ ($s^{-1}$) | $^{conv}K_M$ (g/L) | ε ($s^{-1}$ $g^{-1}$ mL) | $R^2$ | $^{inv}V_{max}/S^*_0$ (nmol $g^{-1}$ $s^{-1}$) | $^{inv}K_M$ (nM) | ε ($s^{-1}$ $g^{-1}$ mL) | $R^2$ |
|---|---|---|---|---|---|---|---|---|---|
| LCC | 0 | 0.29 ± 0.06 | 1.8 ± 1.4 | 161 | 0.9 | 8 ± 1.1 | 86 ± 25 | 94 | 0.87 |
| | 1 | 0.45 ± 0.06 | 2.9 ± 2.3 | 155 | 0.94 | 13.6 ± 1.2 | 94 ± 28 | 144 | 0.92 |
| | 2 | 0.81 ± 0.07 | 4 ± 3.2 | 201 | 0.91 | 24.6 ± 1.3 | 103 ± 30 | 238 | 0.96 |
| | 3 | 1.01 ± 0.09 | 5.2 ± 4.1 | 196 | 0.93 | 30.5 ± 1.4 | 112 ± 33 | 272 | 0.95 |
| | 6 | 1.46 ± 0.19 | 8.5 ± 6.7 | 172 | 0.92 | 37.3 ± 2.1 | 139 ± 40 | 268 | 0.92 |
| | 10 | 0.71 ± 0.15 | 12.9 ± 10.2 | 55 | 0.81 | 9.1 ± 1.4 | 174 ± 51 | 52 | 0.97 |
| | 15 | 0.5 ± 0.15 | 18.5 ± 14.6 | 27 | 0.75 | 0.9 ± 1.2 | 219 ± 64 | 4 | 0.11 |
| | 20 | 0.22 ± 0.12 | 24 ± 19 | 9 | 0.7 | 0.8 ± 1.3 | 263 ± 77 | 3 | 0.12 |
| Global | | | | | 0.95 | | | | 0.97 |
| | CTAB (μM) | $^{conv}V_{max}/E_0$ ($s^{-1}$) | $^{conv}K_M$ (g/L) | ε ($s^{-1}$ $g^{-1}$ mL) | $R^2$ | $^{inv}V_{max}/S^*_0$ (nmol $g^{-1}$ $s^{-1}$) | $^{inv}K_M$ (nM) | ε ($s^{-1}$ $g^{-1}$ mL) | $R^2$ |
| TfC | 0 | 0.04 ± 0.01 | 1 ± 1.2 | 41 | 0.59 | 1.2 ± 0.2 | 77 ± 15 | 15 | 0.95 |
| | 1 | 0.09 ± 0.01 | 1.7 ± 2.1 | 53 | 0.94 | 1.9 ± 0.2 | 80 ± 16 | 23 | 0.97 |
| | 3 | 0.21 ± 0.02 | 3.2 ± 3.9 | 66 | 0.97 | 3.6 ± 0.2 | 86 ± 17 | 42 | 0.96 |
| | 6 | 0.36 ± 0.03 | 5.4 ± 6.7 | 65 | 0.91 | 5.4 ± 0.2 | 95 ± 19 | 56 | 0.97 |
| | 10 | 0.43 ± 0.04 | 8.4 ± 10.4 | 51 | 0.96 | 6.5 ± 0.2 | 107 ± 21 | 61 | 0.95 |
| | 20 | 0.69 ± 0.09 | 15.9 ± 19.5 | 43 | 0.98 | 8 ± 0.3 | 138 ± 27 | 58 | 0.98 |
| | 40 | 0.62 ± 0.12 | 30.8 ± 37.9 | 20 | 0.95 | 4.4 ± 0.3 | 198 ± 39 | 22 | 0.99 |
| | 60 | 0.41 ± 0.09 | 45.8 ± 56.3 | 9 | 0.9 | 2.5 ± 0.3 | 258 ± 51 | 10 | 0.98 |
| Global | | | | | 0.97 | | | | 0.98 |

**Table S1B. Parameters from local non-linear regression.**

| Enzyme | CTAB (μM) | $^{conv}V_{max}/E_0$ ($s^{-1}$) | $^{conv}K_M$ (g/L) | ε ($s^{-1}$ $g^{-1}$ mL) | $R^2$ | $^{inv}V_{max}/S^*_0$ (nmol $g^{-1}$ $s^{-1}$) | $^{inv}K_M$ (nM) | ε ($s^{-1}$ $g^{-1}$ mL) | $R^2$ |
|---|---|---|---|---|---|---|---|---|---|
| LCC | 0 | 0.39 ± 0.05 | 4.9 ± 1.6 | 80 | 0.97 | 8.3 ± 1.3 | 100 ± 54 | 83 | 0.87 |
| | 1 | 0.45 ± 0.06 | 2.9 ± 1.3 | 156 | 0.94 | 13.6 ± 1.6 | 93 ± 39 | 145 | 0.92 |
| | 2 | 0.69 ± 0.07 | 2.2 ± 0.9 | 319 | 0.94 | 24.4 ± 1.9 | 100 ± 28 | 243 | 0.96 |
| | 3 | 1.23 ± 0.27 | 8.3 ± 3.9 | 149 | 0.94 | 30.3 ± 2.8 | 110 ± 35 | 277 | 0.95 |
| | 6 | 1.21 ± 0.23 | 5.4 ± 2.6 | 224 | 0.93 | 37.5 ± 5.1 | 143 ± 60 | 263 | 0.92 |
| | 10 | - | - | 30 | 0.94 | 8.8 ± 0.8 | 154 ± 40 | 57 | 0.97 |
| | 15 | - | - | 17 | 0.89 | 0.5 ± 0.1 | 11 ± 15 | 47 | 0.68 |
| | 20 | - | - | 6 | 0.81 | 0.5 ± 0 | 23 ± 12 | 21 | 0.89 |
| | CTAB (μM) | $^{conv}V_{max}/E_0$ ($s^{-1}$) | $^{conv}K_M$ (g/L) | ε ($s^{-1}$ $g^{-1}$ mL) | $R^2$ | $^{inv}V_{max}/S^*_0$ (nmol $g^{-1}$ $s^{-1}$) | $^{inv}K_M$ (nM) | ε ($s^{-1}$ $g^{-1}$ mL) | $R^2$ |
| TfC | 0 | 0.15 ± 0.05 | 26.9 ± 13.8 | 6 | 0.98 | 1.1 ± 0.1 | 65 ± 18 | 17 | 0.96 |
| | 1 | 0.11 ± 0.01 | 3.9 ± 0.7 | 28 | 0.99 | 1.8 ± 0.1 | 78 ± 18 | 24 | 0.97 |
| | 3 | 0.21 ± 0.02 | 3.1 ± 0.9 | 67 | 0.97 | 3.5 ± 0.2 | 75 ± 20 | 47 | 0.96 |
| | 6 | 0.29 ± 0.04 | 3 ± 1.3 | 99 | 0.94 | 5.4 ± 0.4 | 101 ± 26 | 54 | 0.97 |
| | 10 | 0.44 ± 0.09 | 8.7 ± 3.5 | 50 | 0.96 | 6.6 ± 0.6 | 113 ± 35 | 58 | 0.95 |
| | 20 | 0.81 ± 0.23 | 20.7 ± 8.9 | 39 | 0.98 | 7.9 ± 0.6 | 134 ± 30 | 59 | 0.98 |
| | 40 | - | - | 15 | 0.99 | 4.4 ± 0.2 | 203 ± 27 | 22 | 0.99 |
| | 60 | - | - | 7 | 0.94 | 2.4 ± 0.2 | 225 ± 46 | 11 | 0.98 |

### 2. Control experiments with high concentration of CTAB

In order to investigate any denaturing effect of CTAB on the enzymes, we exposed LCC and TfC to high concentrations of CTAB during 2 hours, prior to activity measurements on the soluble *p*NP-Bu substrate. The results (Figure S3) clearly showed that none of the enzymes were catalytically impaired after the incubation with CTAB.

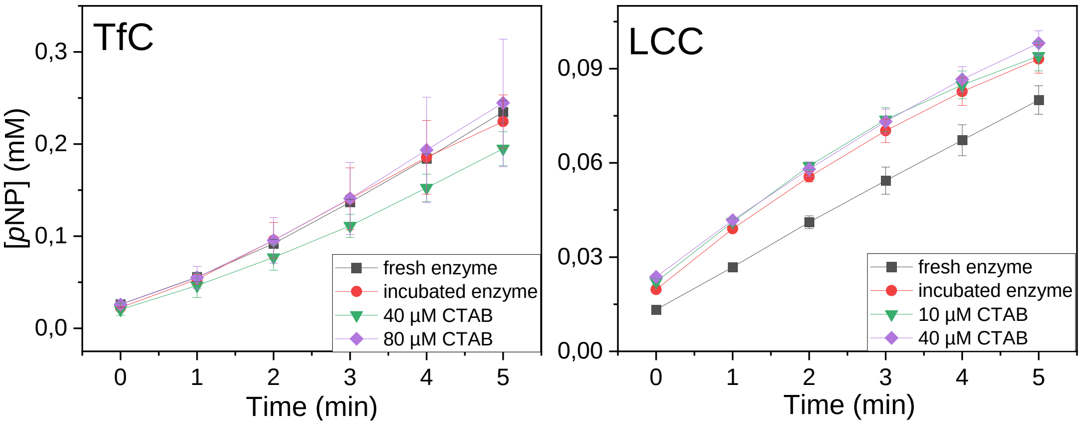


*Figure S3. Progress curves showing the enzymatic hydrolysis of pNP-Bu by TfC (left panel) and LCC (right panel) after being incubated for two hours at 50 °C, alone or together with CTAB. Prior to pNP-Bu hydrolysis, the incubated enzyme reactions were diluted 1000-fold, in order to remove any substantial amount of surfactant that could influence the reaction. A control reaction where enzymes were taken fresh from the fridge was also included. The amount of surfactant was chosen based on concentrations that were contributing to a decrease in activity on the insoluble PET substrate (see Figures 2 and 3 in the main text). The release of pNP was monitored over 5 minutes at 25 °C. Data represent mean values from duplicate experiments, and error bars indicate the spread.*

### 3. Initial activity measurements with increasing amount of CTAB

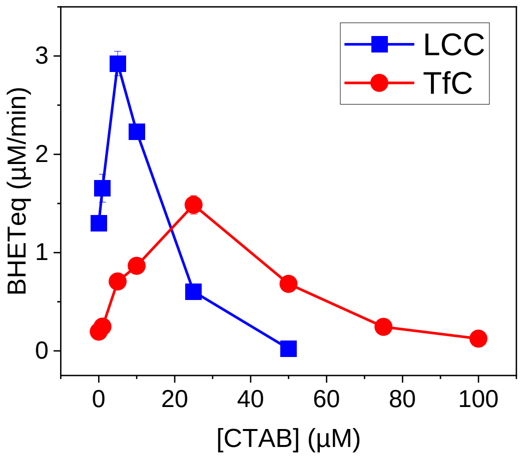


*Figure S4. PET hydrolase activity (quantified as BHETeq/min) at increasing concentrations of CTAB (0-100 µM). Reactions were incubated with 20 g/L PET, 0.1 µM enzyme at 50 °C over 30 min (LCC) or 2 hours (TfC). Experiments were conducted in duplicates and standard errors represent the spread. Both enzymes demonstrate increasing activities up to a certain concentration of surfactant, where the activity starts to level off.*
